# A framework evaluating the utility of multi-gene, multi-disease population-based panel testing that accounts for uncertainty in penetrance estimates

**DOI:** 10.1101/2022.08.10.503415

**Authors:** Jane W. Liang, Kurt D. Christensen, Robert C. Green, Peter Kraft

## Abstract

Panel germline testing allows for efficient detection of deleterious variants for multiple conditions, but the benefits and harms of identifying these variants are not always well-understood. We present a multi-gene, multi-disease aggregate utility formula that allows the user to consider adding or removing each gene in a panel based on variant frequency; estimated penetrances; and subjective disutilities for testing positive but not developing the disease and testing negative but developing the disease. We provide credible intervals for utility that reflect uncertainty in penetrance estimates. Rare, highly-penetrant deleterious variants tend to contribute positive net utilities for a wide variety of user-specified disutilities, even when accounting for parameter estimation uncertainty. However, the clinical utility of deleterious variants with moderate, uncertain penetrance depends more on assumed disutilities. The decision to include a gene on a panel depends on variant frequency, penetrance, and subjective utilities, and should account for uncertainties around these factors.

## Introduction

Genetic screening for deleterious variants (DVs) in genes associated with monogenic hereditary conditions (typically pathogenic and likely pathogenic variants) can be a valuable component of risk management for opportunistic and population-based genomic screening^1,2^. Testing results can prompt heightened surveillance, prophylactic surgery, and other measures to enhance prevention or treatment. Technological advances such as next-generation sequencing have made simultaneous testing of multiple genes cheaper and more accurate than ever before^3,4^. Panel studies have led to many clinically significant findings that would have been missed by single-gene or single-syndrome testing^3–6^. However, the clinical utility of such comprehensive panel germline testing may not be universally appropriate for all contexts.

For some genes and diseases, published guidelines provide best practices about actions to take when deleterious variants are identified in the context of diagnostic testing. For other genes and settings (e.g. secondary findings or population screening), there is a lack of consensus on whether screening itself, or subsequent interventions based upon screening, should be recommended, often because the disease penetrance (the probability that carriers of deleterious variants will develop disease) is low or unknown. For example, while penetrance has been estimated through families with strong family histories, the penetrance estimates in population screening may still be uncertain^7^. If the benefits, risks, and guidelines are unclear for these genes, it could be harmful rather than beneficial to include them in a testing panel. Instead of mitigating risk and improving outcomes, testing may lead to unnecessary surveillance and overtreatment.

We consider a scenario in which the goal is to determine which genes should be included in a panel as part of non-diagnostic screening for a fixed group of diseases. This setting occurs at the stage when a panel is being built and is distinct from the problem of developing a clinician/patient-facing risk tool. While our approach is readily generalizable to other contexts, such as asymptomatic high-risk settings and the incorporation of variants of uncertain significance, we will focus on population-based screening for deleterious variants in asymptomatic individuals as our motivating context. We propose an aggregate utility function that incorporates quantitative measures of genetic and disease characteristics (carrier prevalences and disease penetrances) and utility benefits and harms (harms and costs are sometimes termed “disutility”) for multiple diseases and germline tests. Positive utilities could include identifying individuals at high risk for disease who would benefit from intervention, who would remain unrecognized in the absence of testing. Disutilities could include anxiety or false reassurance in response to test results, unnecessary surveillance, and overtreatment. Utilities and disutilities can be individualized for specific diseases and tests, as well as patient and clinician concerns.

This approach generates a single net utility across all genes proposed for inclusion in a panel, but our construction also allows for the evaluation of each disease and gene combination on its own merits. We note that our notion of net utility and (dis)utilities is distinct from the *health utility* that is frequently used within decision science for valuing a disease state with respect to death and perfect health. This health utility is typically a value between 0 to 1, but our net utility may take on any positive or negative value, with positive net utilities indicating that it is beneficial to include the gene(s) and negative net utilities indicating that it is harmful.

Additionally, we incorporate credible intervals for disease penetrances that reflect our confidence in available penetrance estimates and propagate this uncertainty into the net utility calculation. Uncertainty may be due to lack of sufficient data to estimate prevalences or penetrances for certain DVs and diseases, as well as ancestral populations or other subgroups of interest. For sufficiently large penetrances, the net utility may provide evidence in favor of keeping the test even when the penetrance estimate is unreliable. For low or moderate penetrances, the net utility may point toward removing the gene from the panel or needing to improve the reliability of the penetrance estimate. This utility approach can be used to help formalize the decision-making process when designing a gene panel. Some approaches that take uncertainty in risk estimates into account for other domains include Berry and Parmigiani^8^ and Ding et al.^9^ The former considers quantifying uncertainty for decision analysis of testing for BRCA1 and BRCA2 mutations, while the latter applies Bayesian methods to estimate the variance of an individual’s polygenic risk score.

Some related methodology has been proposed for addressing how to select genes for panel germline testing, as well as the broader question of how to best leverage modern sequencing technology for clinical use. Most identify clinical interpretation and actionability as critical considerations; many focus on diagnostic applications, while we consider non-diagnostic applications (e.g. risk-stratified screening recommendations). Hall et al.^10^ give an overview of the benefits and challenges in gene panel testing for inherited cancer risk assessment, highlighting ambiguous clinical utility as a potential disadvantage. Because genetic testing results may provide uncertainty rather than information for managing cancer risk, the authors recommend testing be used alongside professional consultation. Xue et al.^11^ modify Shashi et al.^12^‘s testing algorithm for evaluating which molecular diagnostic tool (single-gene tests, gene panels, or exome sequencing) to use for diagnostic yield, depending on the clinical setting. Finally, Mazzarotto et al.^13^ develop a “diagnostic effectiveness” score for determining genes to include for hypertrophic cardiomyopathy genetic testing panels, based on variant classification and penetrance. They identify new genes to screen for but also suggest that panels beyond a limited size provide limited additional sensitivity.

An illustration of our approach to germline testing for deleterious variants on five genes (*ATM, BRCA1, BRCA2, CHEK2*, and *PALB2*) associated with increased risk of developing breast cancer follows in the Results section, along with a broad exploration of the effects of varying parameter inputs (true penetrance, uncertainty in penetrance estimates, variant frequencies, relative (dis)utilities). The Methods section details our formulation of general expressions for our proposed multi-gene, multi-disease aggregate utility. Our method is implemented in an R Shiny app, freely accessible at https://janewliang.shinyapps.io/agg_utility, where users can enter parameter estimates and uncertainties for calculating their own net utilities.

## Results

### Female breast cancer application

We first consider a specific application for the aggregate utility approach that incorporates panel germline testing for *ATM, BRCA1, BRCA2, CHEK2*, and *PALB2* as part of risk assessment for female breast cancer, for hypothetical screening of a woman without a previous breast cancer diagnosis or breast cancer family history. These five genes are commonly included in risk panels for hereditary breast cancer. We chose this example for its familiarity and relevance, as well as the availability of empirical estimates of the lifetime risk of breast cancer in women for carriers of deleterious variants in these genes and their relative precisions. We stress that these results are largely presented for illustrative purposes. In particular, although our understanding of the absolute and relative uncertainty in the penetrance estimates for these genes is changing as more data become available in more diverse populations, the lifetime risk for carriers of DVs in these genes is relatively well known—the uncertainty in penetrances estimates for other diseases and other genes is often much greater^14,15^. Users are free to input their own prevalence and penetrance estimates, as well as the uncertainty in the penetrance estimates, into our R Shiny app to calculate their impact on likely net utility.

Deleterious variants in these genes have all been linked to breast cancer, but some are better studied than others. *BRCA1* and *BRCA2* DVs are highly penetrant with widely adopted guidelines for enhanced screening and other clinical interventions^16–19^. While DVs of *ATM, CHEK2*, and *PALB2* have been linked to breast cancer, the additional risk conferred is not as well understood^20–23^, especially among individuals with non-European ancestries. In this section, quantities (prevalences and penetrances) involving *ATM, BRCA1, BRCA2*, and *PALB2* are estimated for any deleterious variant in the given gene; for *CHEK2*, quantities are for the 1100delC variant only.

The carrier prevalences for DVs on *BRCA1* (0.00058) and *BRCA2* (0.00068) are calculated based on allele frequency estimates reported in Antoniou et al.^24^ (see also Dullens et al.^25^; Krassuski et al.^26^). Those for *ATM* (0.0019), *CHEK2* (0.0026), and *PALB2* (0.00057) are calculated based on allele frequencies reported in Lee et al.^27^ Cumulative lifetime penetrance estimates for female breast cancer are taken from the literature review performed by the All Syndromes Known to Man Evaluator^28–32^: 0.35 (*ATM*), 0.73 (*BRCA1*), 0.72 (*BRCA2*), 0.19 (*CHEK2*), and 0.38 (*PALB2*). This is the genotype-specific probability of developing breast cancer among females, prior to dying. In this illustration, we use parameters estimated from large meta-analyses, largely of studies from the USA, UK, Australia, or countries in Western Europe. Estimates for specific ancestries and subgroups may be used instead, to better reflect a different population of interest

To reflect our greater confidence in the penetrance estimates for *BRCA1* and *BRCA2*, we use a precision of 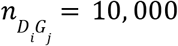 to specify the parameters in their uncertainty distributions. 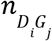 parameterizes the uncertainty in estimating the penetrance for disease *i* associated with deleterious variants in gene *j*; see Methods. Intuitively, the penetrance’s uncertainty distribution can be thought of as a posterior distribution from a trial with 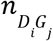 carriers, where larger values of 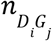 correspond to a greater degree of perceived certainty for the estimate. For *ATM, CHEK2*, and *PALB2*, we specify 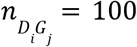, i.e. a smaller trial size of 100. (We chose these values to illustrate the impact of uncertainty on net utility calculations. They should not be taken as indicative of the absolute or relative strength of the available data on the penetrance of DVs in these genes.) Supplementary Figure 1 plots the uncertainty distributions of the five lifetime penetrance estimates. The wider spread for *ATM, CHEK2*, and *PALB2* reflects greater uncertainty. Supplementary Table 1 summarizes the quantiles for these uncertainty distributions at 2.5%, 5%, 10%, 50%, 90%, 95%, and 97.5%.

We report the individual gene net utilities and aggregate utility for a multigene breast cancer panel testing for deleterious variants in all five of these genes. Individual gene utility is the net change in utility from including a gene in the panel relative to not screening for deleterious variants in that gene; the aggregate utility for a set of genes (and diseases) is the sum of individual gene (and disease) utilities (see Methods). The aggregate net utility Δ is a function of the frequency of deleterious variants, lifetime penetrances, and relative disutilities 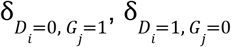, and 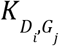 (indexed over gene *i* and disease *j*). Further detail is provided in Methods, but in brief, 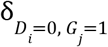 denotes the disutility for an individual who tests positive for the gene, but does not develop the phenotypic features of the associated disease (abbreviated G+D-); 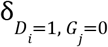 denotes the disutility for an individual who tests negative for the gene, but does develop phenotypic features of the associated disease (abbreviated G-D+); and 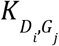 is the disutility of screening itself. Because hereditary predisposition for breast cancer drives only a portion of cases, most individuals in the general population who develop breast cancer over their lifetimes would fall in the G-D+ group. Since the deleterious variants considered are rare, G+D-individuals (those with hereditary cancer predisposition due to these genes) represent a small subgroup of those who never develop breast cancer. In general, the disutilities encompass a broad range of financial and non-financial harms and can vary across disease *i* and tested gene *j*. For ease of presentation, we assume that disutilities are the same across all five tests (denoted as δ_*D*=0, *G*=1_, δ_*D*=1, *G*=0_, and *K*), that *K*=0, and (without loss of generality) that δ_*D*=0, *G*=1_ = 1. We allow the utility ratio δ_*D*=1, *G*=0_ /δ_*D*=0, *G*=1_ to vary from 0.1 to 10 in increments of log_10_ (0. 1) (see Methods). Figure 1 plots the net utilities against this utility ratio for each of the individual genes, as well as the aggregate Δ for all five genes. Supplementary Figure 2 depicts the same curves on a log-transformed x-axis, to help illustrate the behavior for small values. Supplementary Figure 3 presents a supplemental analysis where the disutilities for BRCA and BRCA2 differ from those specified for ATM, CHEK2, and PALB2.

**Figure 1:**
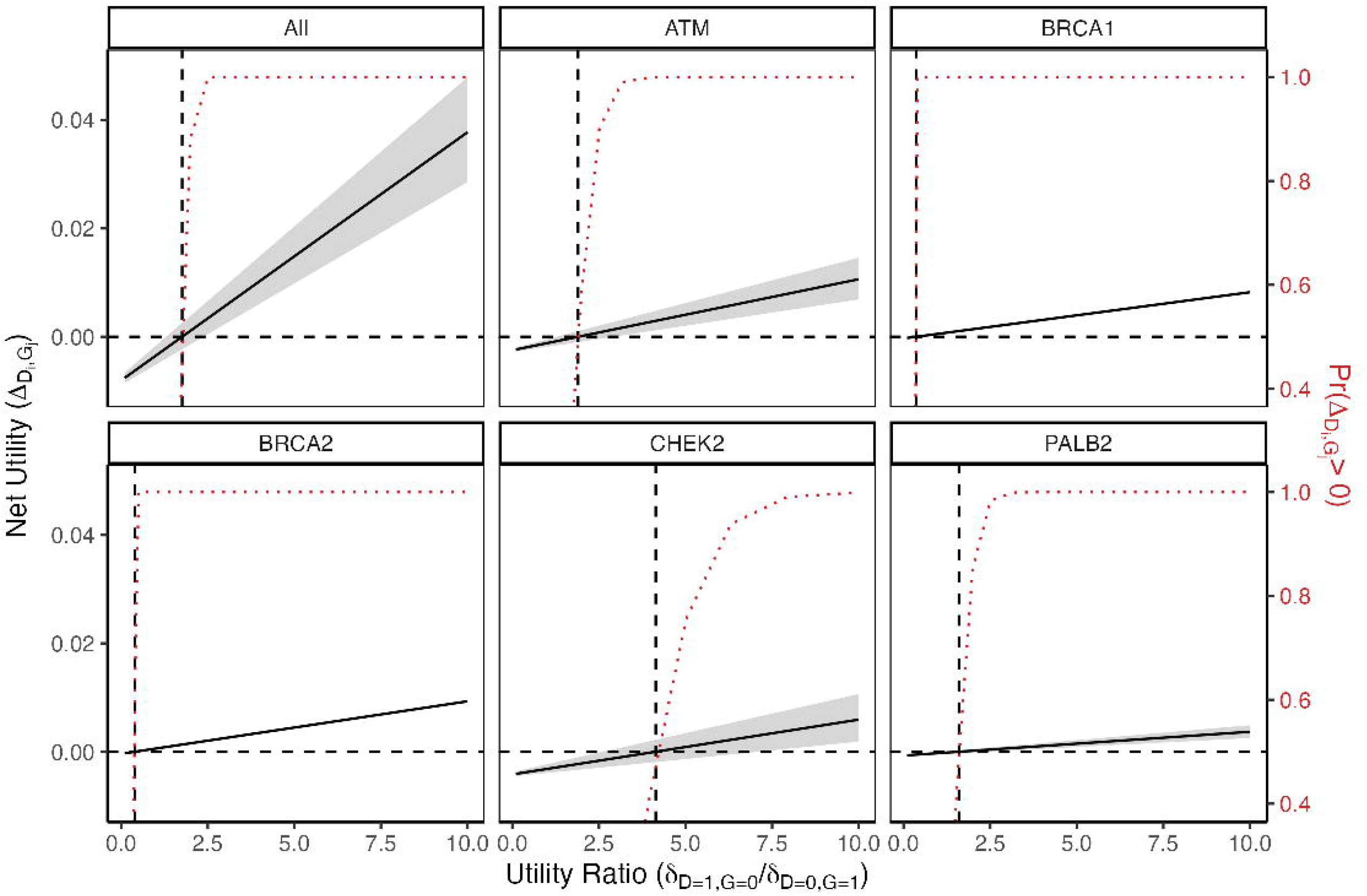
Net utilities from the female breast cancer application plotted against the ratio δ_*D*=1, *G*=0_ /δ_*D*=0, *G*=1_ for each of the individual genes, as well as the aggregate Δ for all five genes. δ_*D*=1, *G*=0_ is allowed to vary from 0.1 to 10 in increments of log_10_(0. 1), while fixing *K*=0 and δ_*D*=0, *G*=1_ = 1. 95% credible intervals are shaded in gray and represent uncertainty contributed by the penetrance estimates taken from a literature review (see text). Dashed reference lines are drawn to indicate the value of the ratio at which the net utility changes from negative to positive. The red dotted curve represents the probability of a positive net utility, with respect to the uncertainty distribution, at each value of δ_*D*=1, *G*=0_ /δ_*D*=0, *G*=1_.

As expected, the credible intervals for the *BRCA1* and *BRCA2* net utilities are very narrow and the credible intervals for *ATM, CHEK2*, and *PALB2* are wider, reflecting the widths of the credible intervals in their uncertainty penetrance distributions. The aggregate utility has the widest credible intervals of all, because it incorporates uncertainty from all five penetrance estimates.

We can consider interpreting the results in terms of utility thresholds, defined as the ratio of 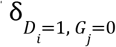 (disutility of G-D+) to δ_*D*=0, *G*=1_ (disutility of G+D-) such that the net utility is 0 (Table 1). Additional detail, including derivations, can be found in the Methods, but in general, a gene with a higher utility threshold (>1) can be interpreted as having a more limited range of subjective inputs 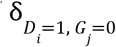 and δ_*D*=0, *G*=1_ where it would still be beneficial to include it in the panel. These threshold values are quite low for *BRCA1* (0.37) and *BRCA2* (0.40), so even in a scenario where one is highly concerned with avoiding G+D-results, there is a wide range of possible disutilities that can be specified to result in a positive net utility. The net utility for testing these two genes is positive, except for some extreme cases when δ_*D*=1, *G*=0_ is very low compared to δ_*D*=0, *G*=1_. The curves for the probability of the net utility being positive resemble step functions, with the jump from being 0% positive to 100% positive occurring at a sharp, early point.

**Table 1:** Female breast cancer application utility thresholds 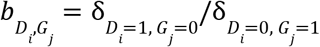 at which the net utility is 0 for the five individual genes and overall (“All”). Lower threshold values (below 1) indicate that a positive net utility can be achieved with a lower G-D+ disutility, relative to the G+D-disutility. The upper and lower bounds for a 95% credible interval are also reported.

|  | Estimate | 0.025 | 0.975 |
| --- | --- | --- | --- |
| **All** | 1.8 | 1.3 | 2.4 |
| **ATM** | 1.9 | 1.3 | 2.9 |
| **BRCA1** | 0.37 | 0.35 | 0.38 |
| **BRCA2** | 0.40 | 0.38 | 0.41 |
| **CHEK2** | 4.1 | 2.6 | 7.1 |
| **PALB2** | 1.6 | 1.08 | 2.42 |

In contrast, the less-penetrant genes have utility thresholds above 1: *ATM* (1.9), *CHEK2* (4.1), and *PALB2* (1.6). In these cases, the G-D+ disutility needs to outweigh the G+D-disutility in order for it to be beneficial to keep the gene in the panel, sometimes by a considerable amount. Because of the greater uncertainty in the penetrance estimates, the lower bound of the utility threshold credible interval (Table 1) is noticeably even less favorable. The probability curve for observing a positive net utility also bends toward 100% at a much more gradual incline. The aggregate utility threshold is somewhere intermediate (1.8), balancing between the larger and smaller effect sizes, as is the shape of the probability curve.

Heatmaps of the individual and aggregate net utilities while holding *K*=0 and varying δ_*D*=0, *G*=1_ and δ_*D*=1, *G*=0_ from 1 to 100 in increments of 5 are depicted in Figure 2. Similar heatmaps for the probability of a positive net utility and the fifth percentile net utility are shown in Supplementary Figures 4 and 5. Utility thresholds based on the original penetrance estimates are drawn as solid black lines on all three sets of heatmaps. The dashed black reference lines have intercept 0 and slope 1, and correspond to cases where G+D- and G-D+ have equal disutilities. In Supplementary Figure 5, the additional solid blue lines indicate the utility thresholds based on the fifth percentiles of the penetrances’ uncertainty distributions.

**Figure 2:**
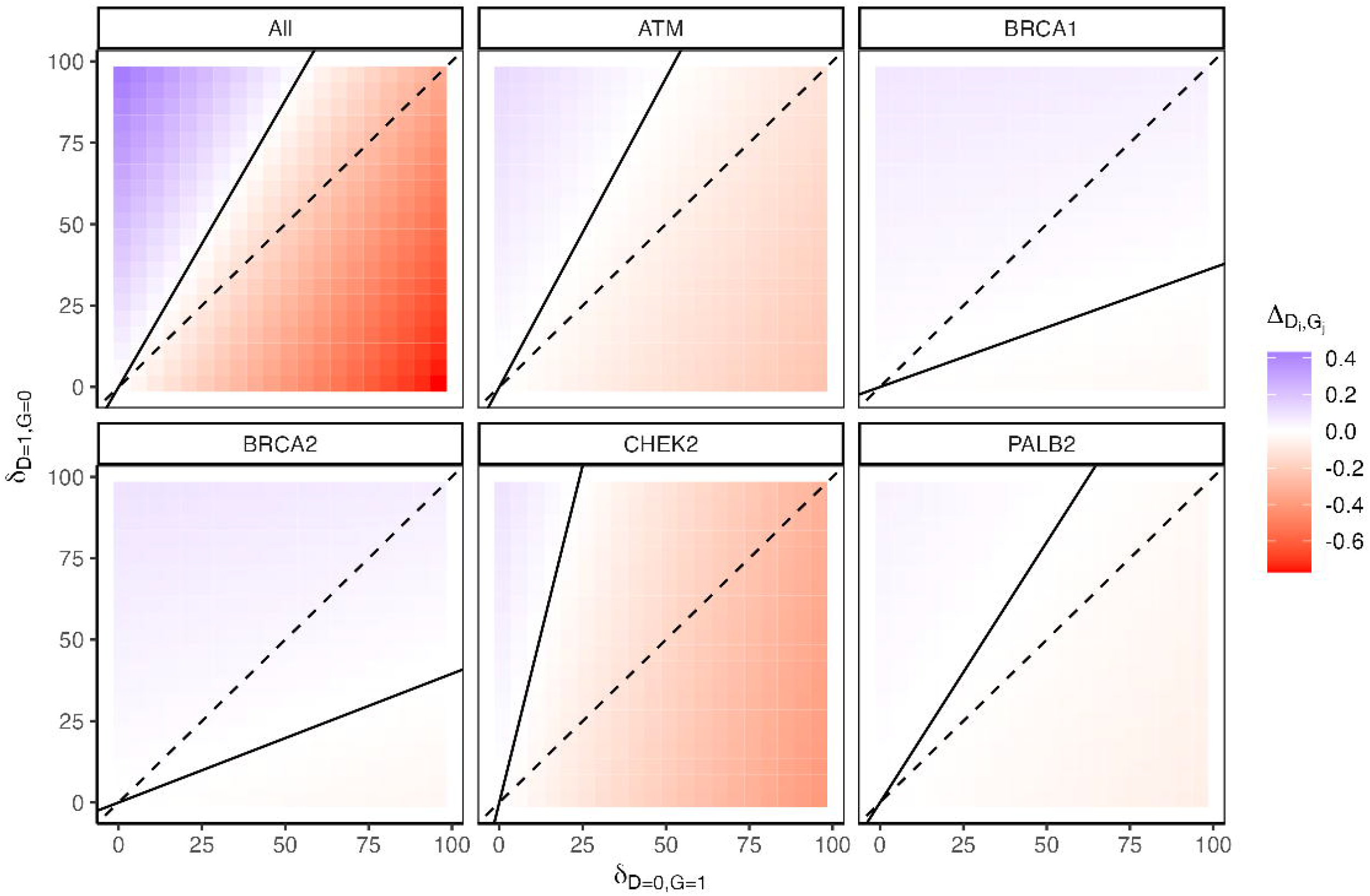
Heatmaps of the five individual female breast cancer net utilities and the aggregate utility (“All”) while holding *K*=0 and varying δ_*D*=0, *G*=1_ and δ_*D*=1, *G*=0_ from 1 to 100 in increments of 5. Positive net utilities are shaded blue and negative utilities are shaded red. Utility thresholds based on the penetrance estimates are drawn as solid black lines. The dashed black lines have intercept 0 and slope 1, and correspond to cases where the individual testing positive for the gene but not developing the disease and vice versa have equal disutilities.

These heatmaps offer an alternative visualization as well as some additional insight on the behavior of the utilities under different G+D- and G-D+ disutility conditions. In Figure 2, the net utilities for *BRCA1* and *BRCA2* are positive (blue) for a much broader range of δ_*D*=0, *G*=1_ and δ_*D*=1, *G*=0_ values compared to the net utilities for the other genes, in concordance with the utility threshold discussion for Figure 1.

The utility threshold reference lines in the heatmaps for the probability of a positive net utility (Supplementary Figure 4) track with the regions where the probability of positive net utility transitions from 0 (white) to 1 (dark blue). The sharp transitions for *BRCA1* and *BRCA2* reflect their tight credible intervals, and the more gradual transitions for *ATM, CHEK2*, and *PALB2* reflect their wider credible intervals. The heatmaps for the fifth percentiles of the net utilities (Supplementary Figure 5) closely resemble the heatmaps for the net utilities based on the original penetrance estimates. The utility thresholds for *BRCA1* and *BRCA2* are quite similar; there is more variability in the utility thresholds for *ATM, CHEK2*, and *PALB2*. Again, this reflects the wider credible intervals and uncertainty distributions for these genes. Under a “near-worst case scenario” interpretation (i.e. basing decisions about potential utility on the lowest 5th percentile of the utility distribution), *ATM, CHEK2*, and *PALB2* require the specification of an even larger G-D+ disutilities relative to their G+D-disutilities in order to result in positive net utilities for testing.

### Net utility behavior as parameters vary

In order to explore the properties of our proposed net utility expression across a wider range of scenarios, we varied the parameters influencing the individual net utility for disease *i* and gene *j*, 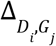, as follows. Let*D* _*i*_ = {0, 1} be the indicator for developing disease *i* and *G*_*j*_ = {0, 1} be the indicator for testing positive for carrying a deleterious variant on gene *j* (see Methods).

- G-D+ disutility 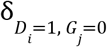: Ranging from 0.1 to 10 in increments of log_10_(0. 1)
- Cumulative lifetime disease penetrance *Pr*(*D*_*i*_ = 1 | *G*_*j*_ = 1): {0. 2, 0. 4, 0. 6, 0. 8, 0. 99}
- Carrier prevalence *Pr*(*G*_*j*_ = 1): {0. 001, 0. 002, 0. 003, 0. 004}
- Precision 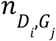 used to specify parameters in the uncertainty distribution: {10, 100, 1000, 10000}

The chosen penetrance and carrier prevalence values reflect those seen in clinical practice. Lifetime penetrances vary between 0.195 and 0.732 and carrier prevalences vary between 0.00114 and 0.00519 in the female breast cancer application. The ClinGen actionability reports^33^ frequently list disease risks with broad ranges of possible values. For example, carriers of *STK11* deleterious variants have a 38-66% estimated risk of developing gastrointestinal cancer by age 60-70 and a 13-18% risk for gynecological cancer^34^. The penetrance of developing dopa-responsive dystonia among *GCH1* carriers is 87-100% for females and 35-55% for males^35^. Carriers of *MLH1, MSH2, MSH6* or *PMS2* have a 25-70% cumulative risk for colorectal cancer by age 70 and 30-70% for endometrial cancer. Penetrances by age 70 for other cancers are generally lower in effect size and narrower in range of estimated values, including 1-9% for gastric, 2-16% for bladder, 6-14% for ovarian, 9-30% for prostate, and 5-14% for breast^36^.

As in the previous subsection, we set the test disutility 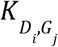 to 0 and the G+D-disutility 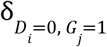 to 1, thereby normalizing the utility ratio to be 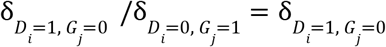 and allowing it to range from 0.1 to 10 in increments of log_10_ (0. 1). Figure 3 plots 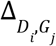 against 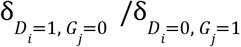 for each combination of parameters, with prevalences varying in the rows and penetrances varying in the columns. (Supplementary Figure 6 depicts the same curves on a log-transformed x-axis, to help illustrate the behavior for small values.) The colored shading represents 95% credible intervals for different values of 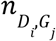. Dashed reference lines are drawn to indicate the utility threshold in each scenario. Table 2 gives these threshold values with a 95% credible interval.

**Table 2:** Utility thresholds 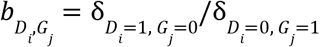 at which the net utility is 0 while varying the G-D+ disutility, disease penetrance, carrier prevalence, and precision parameter. Lower threshold values (below 1) indicate that a positive net utility can be achieved with a lower G-D+ disutility, relative to the G+D-disutility. The upper and lower bounds for a 95% credible interval are also reported.

| Penetrance | Precision | Estimate | 0.025 | 0.975 |
| --- | --- | --- | --- | --- |
| **0.2** | **10** | **4** | 1.1 | 35 |
|  | **100** |  | 2.5 | 6.8 |
|  | **1000** |  | 3.4 | 4.7 |
|  | **10000** |  | 3.8 | 4.2 |
| **0.4** | **10** | **1.5** | 0.43 | 6.3 |
|  | **100** |  | 1.0 | 2.3 |
|  | **1000** |  | 1.3 | 1.7 |
|  | **10000** |  | 1.4 | 1.6 |
| **0.6** | **10** | **0.67** | 0.16 | 2.3 |
|  | **100** |  | 0.44 | 0.99 |
|  | **1000** |  | 0.59 | 0.76 |
|  | **10000** |  | 0.64 | 0.69 |
| **0.8** | **10** | **0.25** | 0.029 | 0.93 |
|  | **100** |  | 0.15 | 0.40 |
|  | **1000** |  | 0.21 | 0.29 |
|  | **10000** |  | 0.24 | 0.26 |
| **0.99** | **10** | **0.01** | 0.000 | 0.11 |
|  | **100** |  | 0.000 | 0.038 |
|  | **1000** |  | 0.005 | 0.017 |
|  | **10000** |  | 0.008 | 0.012 |

**Figure 3:**
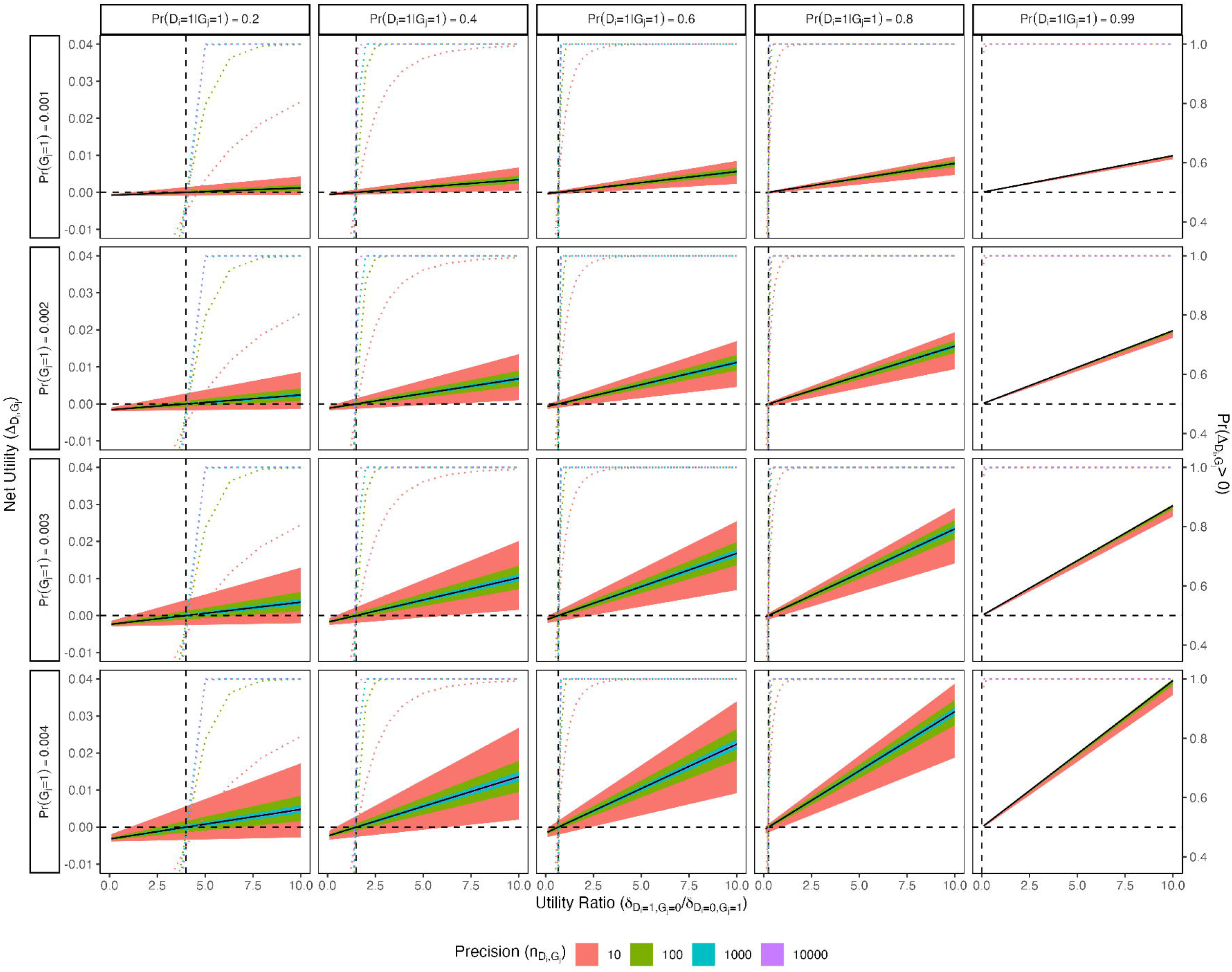
Net utilities for a single gene and disease plotted against the utility ratio 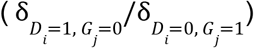, while varying the G-D+ disutility, disease penetrance, carrier prevalence, and precision parameter. Test and G+D-disutilities are fixed at 0 and 1, respectively. The colored shading represents 95% credible intervals for different values of the precision. Dashed reference lines are drawn to indicate the utility threshold in each scenario. The dotted curves represents the probability of a positive net utility, with respect to the uncertainty distribution for each precision level and at each value of 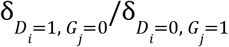.

Overall, higher carrier prevalences and lower penetrances tend to correspond to wider credible intervals for all precision levels. However, if the carrier prevalence is very low, the credible intervals remain consistently narrow even when the penetrance is also very low, and similarly when the disease penetrance is very high in the presence of high prevalence. So the net utilities for rare, highly penetrant genes are more likely to have narrow credible intervals, independent of the amount of confidence we have about the penetrance estimates.

Interestingly, the credible intervals for net utilities at given prevalence and penetrance values grow wider as 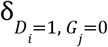 (disutility of a G-D+ result) increases relative to 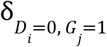 (disutility of a G+D-result). So, even supposing that one undervalues the relative disutility of G+D-, the uncertainty from the distribution of the penetrance allows the decision of which genes to keep in the panel to be less dependent on the exact choice of disutilities. Higher penetrances correspond to lower utility thresholds (since we set 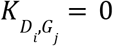, the utility threshold does not depend on prevalence), which makes intuitive sense: high penetrance implies the proportion of carriers who do not develop the disease of interest is small, so interventions with smaller 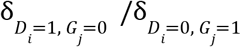 ratios can still have positive net utility.

## Discussion

We have derived net utility expressions to aid in determining which genes add or detract utility from a genomic testing panel in the setting of population-based screening for deleterious variants in asymptomatic individuals. These expressions are functions of carrier prevalence and disease penetrance estimates, as well as user-specified disutilities for G+D-, G-D+, and testing. Our approach is flexible, and allows users to estimate impact in a variety of clinical contexts, from population-level applications to screening of high-risk populations. These expressions may provide a useful framework for determining which genes to include on a custom sequencing panel or which genes to include on a clinical report from whole exome sequencing, for asymptomatic individuals. One goal in this context is to prevent or mitigate the disease of interest by identifying high-risk individuals. The trade-offs in utility will therefore depend partly on the efficacy of the would-be-prescribed interventions for preventing poor health outcomes. We present utility thresholds, the probability of a positive utility, and lower bounds on the net utilities as summary values that can provide additional insight. As an illustration of our approach, we evaluated the net utility of population screening for deleterious variants in five breast cancer predisposition genes, as well as a hypothetical range of disease penetrances, carrier prevalences, and precision values (used for specifying the penetrance’s uncertainty distribution).

Our work provides a needed approach for estimating the incremental utility or disutility of genetic screening for DVs in numerous genes and conditions simultaneously. Published estimates about the clinical benefits of genetic screening to date have focused on conditions with reasonably developed evidence bases^37–40^. Yet, the ability of genomic sequencing to identify genetic variants associated with rare disorders, for which epidemiological evidence is typically limited, has been one of the most promising successes from advances in genetic testing capabilities^41–44^. Moreover, the American College of Medical Genetics and Genomics is integrating an increasing number of conditions into its recommendations for a minimal list for secondary findings disclosure, even when data about the penetrance of DVs in associated genes is limited^45–47^. The approach we have developed allows for better estimation of the benefits and harms of such recommendations, estimates that have been omitted from research to date^40^. Our tool provides a flexible approach that can accommodate varying measures of utility and disutility, including quality-adjusted life years, life years gained or lost, and death rates. Moreover, the tool can be easily tailored to accommodate utility and disutility for a variety of perspectives, from patient outcomes to societal impact^48^.

Our approach does not directly account for potential challenges in curation accuracy for deleterious variants, and we generally do not distinguish between different DVs of the same gene (although our framework is easily modified to have several individual variants or classes of variants). We assume that modern germline testing technology detects DVs with near-perfect sensitivity and specificity. Here, we refer to sensitivity and specificity with respect to sequencing technology, not the problem of classifying variants as benign or deleterious on the basis of clinical sensitivity and specificity. We further assume that the carrier prevalences can be estimated with a high degree of accuracy, such that they do not contribute a significant amount of additional uncertainty to the net utilities. When formulating the net utility expression, we treat untested individuals as being equivalent to those who test negative for the DV(s) in question. In scenarios that deviate considerably from these conditions, we acknowledge that our approach may be of limited use.

Further work can explore challenges when building a utility that incorporates many more genes/variants or genes with unknown parameter estimates/variants of uncertain significance, as well as accounting for the age of the person being tested or measured polygenic risk scores^47,49^. Accounting for the age of the person tested and treating penetrances as age-based distributions would allow us to model more complex relationships between DVs and diseases with incomplete penetrances. We can also conduct a more rigorous exploration of simplifying assumptions that reduce the number of disutility parameters that need to be specified. Nevertheless, our work provides a feasible approach to estimating the clinical benefits or harms of genetic screening. Tools such as ours are critically needed by policymakers and payers as they make decisions about how to regulate and reimburse the current generation of genomic tests.

## Methods

### Utility notation

Suppose that we are interested in risk assessment for some predetermined set of diseases, indexed *i* = 1, …, *I*, and are considering the genes *j* = 1, …, *J* to be included in a panel for germline testing. We define an aggregate utility expression in terms of the following notation:

- *D*_*i*_ = {0, 1} is the indicator for developing disease *i*.
- *G*_*j*_ = {0, 1} is the indicator for testing positive for carrying a deleterious variant on gene *j*.
- 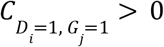 is the utility associated with the scenario where the individual tests positive for carrying a deleterious variant on gene *j* and does develop disease *i* (abbreviated G+D+).
- 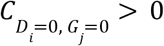 is the utility associated with the scenario where the individual tests negative for carrying a deleterious variant for gene *j* and does not develop disease *i* (abbreviated G-D-).
- 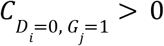 is the utility associated with the scenario where the individual tests positive for carrying a deleterious variant on gene *j* but does not develop disease *i* (abbreviated G+D-). Assume that 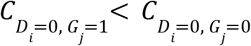.
- 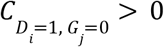 is the utility associated with the scenario where the individual tests negative for carrying a deleterious variant for gene *j* but develops disease *i* (abbreviated G-D+). Assume that 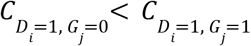.

We emphasize that the scenarios outlined by these definitions capture incomplete penetrance, as opposed to genotyping errors or misclassifying deleterious variants (DVs). In our notation, developing a disease (D+) refers to lifetime development of specific phenotypic features of a condition, and the converse (D-) refers to not developing those features. Let 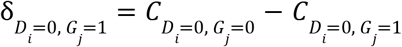 be the disutility associated with testing positive for gene *j* but not developing disease *i* (G+D-) or alternatively the utility benefit of testing negative for gene *j* and not developing disease *i* (G-D-), e.g. the disutility associated with unnecessary surveillance and over-treatment and possible anxiety due to a positive test. Similarly, define 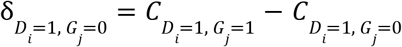 as the disutility associated with testing negative for gene *j* but developing disease *i* (G-D+) or alternatively the utility benefit of testing positive for gene *j* and developing disease *i* (G+D+), e.g. the disutility associated with default screening or preventive interventions relative to more intensive interventions along with false reassurance among those who go on to develop disease. We assume both 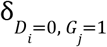 and 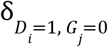 are greater than 0. In other words, we assume that if one does not develop the disease, testing negative for the associated DV leads to more beneficial outcomes, and if one does develop the disease, testing positive leads to more beneficial outcomes. (We do not consider situations where either 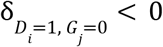 or 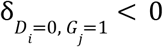, although we note that these may exist; for example, where the G-D+ utility is larger than the G+D+ utility and “the cure is worse than the disease”.) Finally, let 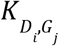 be the disutility (potentially including psychological or physical harms) associated with conducting the test for gene *j* in relation to disease *i*, independent of test results. Then the net utility for disease *i* in the setting where we test for gene *j* is

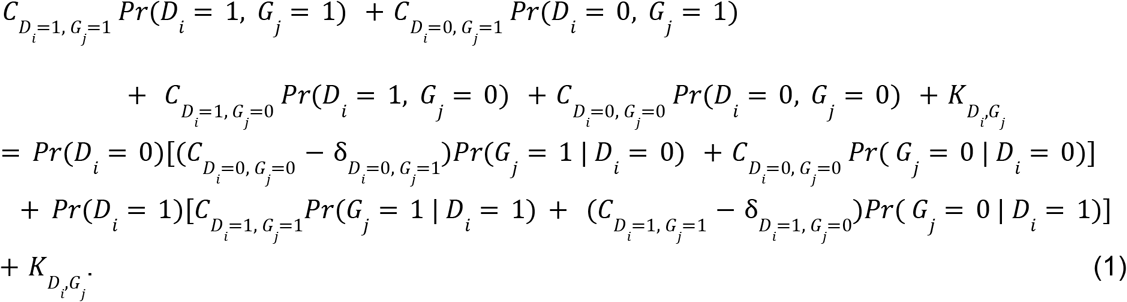

Assuming that the utility associated with developing disease *i* in the absence of testing information for gene *j* is equal to 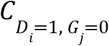, and assuming that the utility for not developing disease *i* in the absence of testing is equal to 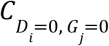, then the net utility for disease *i* in the scenario where we do not test for gene *j* is

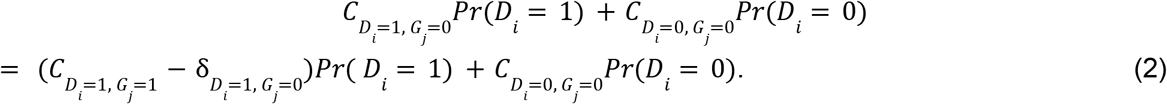

Of interest is the difference in utility for disease *i* when testing vs. not testing for gene *j*, which we define as the difference of Eq. (1) and Eq. (2):

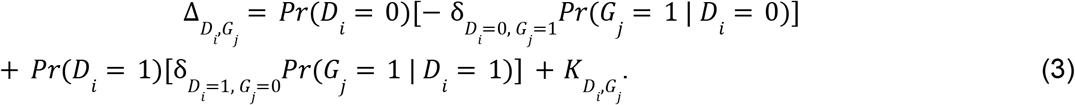

This difference in utility can be re-expressed in terms of *Pr*(*G*_*j*_ = 1), which is the prevalence for DVs of gene *j*, and *Pr*(*D* _*i*_ = 1 | *G*_*j*_ = 1), which is the cumulative lifetime risk or penetrance of developing disease *i* given that one is carries a DV in gene *j* (i.e., the penetrance):

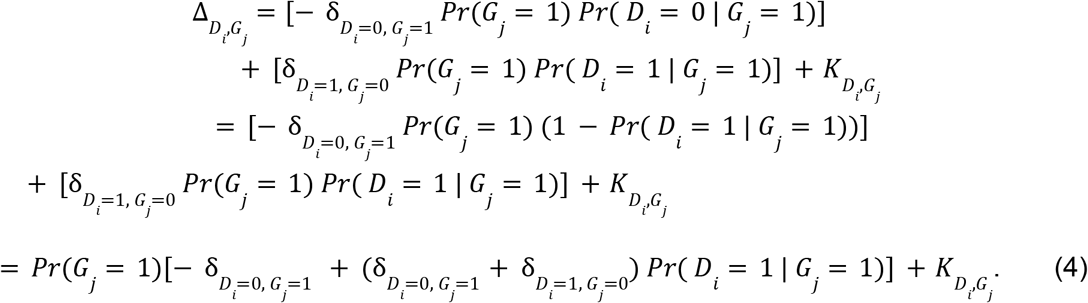

It is beneficial to test for gene *j* when 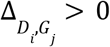, which occurs when the utility for testing is greater than the utility for not testing. For simplicity, we will generally treat testing for a given gene as testing for a particular deleterious variant in the gene, but the framework readily extends to handle variant-specific tests, prevalences, and penetrances.

For multiple diseases (indexed by *i*) and tests (indexed by *j*), the aggregate utility Δ sums over all combinations of *i* and *j*. Doing so requires carrier prevalence and disease penetrance estimates for each gene and disease, as well as the specification of disutilities for testing positive for gene *j* but not developing disease *i* (G+D-), testing negative for gene *j* but developing disease *i* (G-D+), and testing itself. Δ provides a simple summary value while still allowing genes to be evaluated individually:

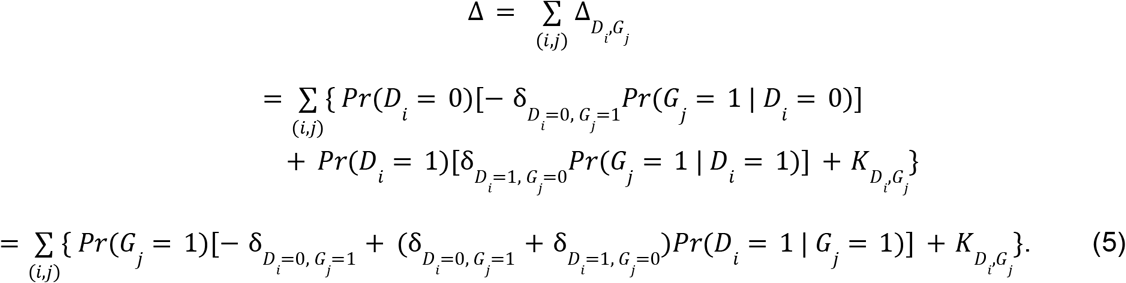

An additional 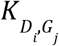 can be included for each (*i, j*) combination (with perhaps less weight given for each additional test), or a single overall disutility for all testing can be used. Since Δ is the sum of the net utilities of particular disease-gene pairs (*i, j*), the decision as to whether or not to include a given test on a multi-gene, multi-disease panel depends only on the individual net utility of that test.

The number of disutility parameters 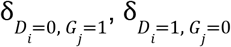, and 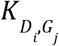 grows as the number of diseases and tests increases, but one can consider simplifications such as assuming the same (dis)utilities across diseases/tests or subgroups of diseases/tests. For example, it may be reasonable to assume that the disutility of each test for an additional gene *j* is negligible. Specification of these disutilities is largely subjective and should depend on the clinical setting and patient concerns.

### Utility threshold

As a general guide for interpretation, if the user fixes the value of 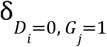, they can conceive of the value of 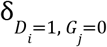 as being a relative weight for the disutility of G-D+ vs the disutility of G+D-. More formally, for an individual test for disease *i* and gene *j*, a utility threshold can be defined as the value 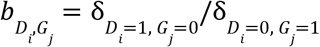 for which 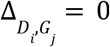:

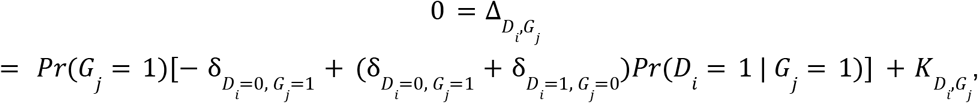

which implies

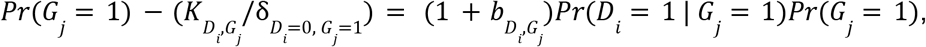

so

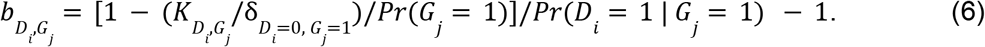

Note that when 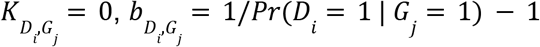 and depends only on the penetrance *Pr*(*D*_*i*_ = 1 | *G*_*j*_ = 1). If the ratio of the disutility of G-D+ to the disutility of G+D- is greater than 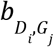, then including gene *j* to test for disease *i* has positive net utility. If the ratio needed to achieve a non-negative utility is unreasonable—e.g. in many settings ascribing a higher disutility to testing positive for the gene but not developing the disease compared to the disutility of testing negative for the gene but developing the disease would be inappropriate—then the test should not be kept as part of the panel. Basing analysis around a threshold ratio allows for an alternative interpretation that does not require upfront specification of the disutilities 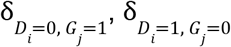, and 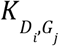.

If one assumes that 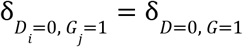 and 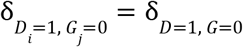 for all values of *i* and *j*, then

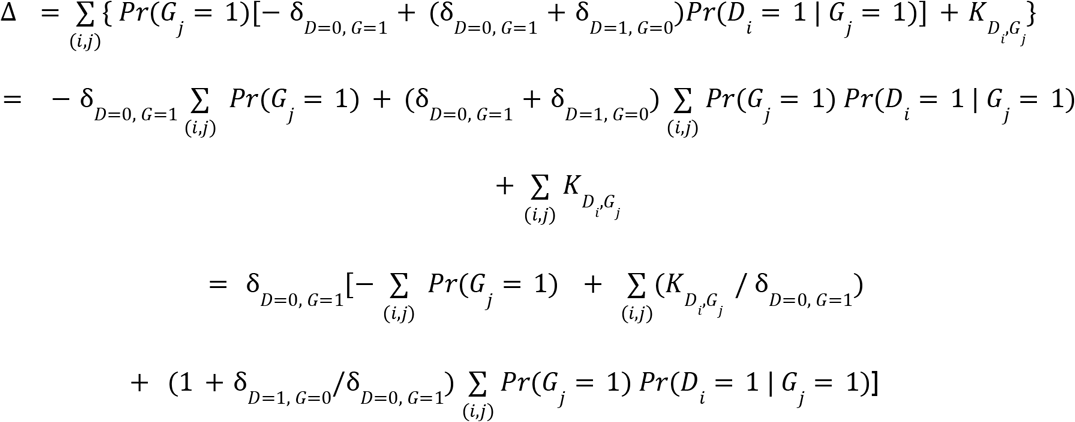

and the threshold *b* = δ_*D*=1, *G*=0_ /δ_*D*=0, *G*=1_ when Δ = 0 can be expressed as

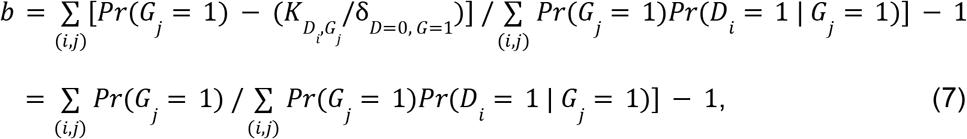

where the last line holds when 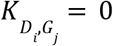 for all *i, j*.

### Uncertainty distribution for disease penetrance

Of additional interest is the incorporation of uncertainty in the penetrance estimates *Pr*(*D*_*i*_= 1 | *G*_*j*_ = 1). Denoting 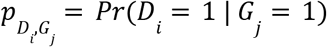 for a given disease *i* and gene *j*, we model the uncertainty in the penetrance 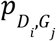 as a beta distribution Beta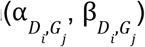. One can motivate the choice of the parameters 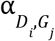 and 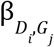 by conceiving of the penetrance’s uncertainty distribution as the posterior distribution from a trial of 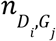 carriers of deleterious variant *j*. Then, set 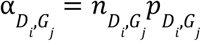 to represent the expected number of cases of disease *i* in the trial and set 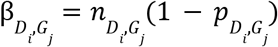 to represent the expected number of individuals who do not develop the disease. Through specification of the precision 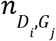, we can express our confidence level in the estimation of 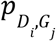, with larger values of 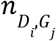 corresponding to a greater degree of certainty about the estimate and smaller ones indicating less confidence.

The uncertainty from 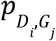 can then be propagated into a distribution and credible interval for the corresponding 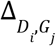 and the aggregate Δ (assuming independence of 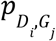 across all *i, j*), as well as additional summary values. We will assume that we are not concerned about incorporating uncertainty from estimating *Pr*(*G*_*j*_ = 1). The probability that the individual net utility 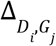 is positive (i.e. adding the test for gene *j* makes an improvement) can be written as

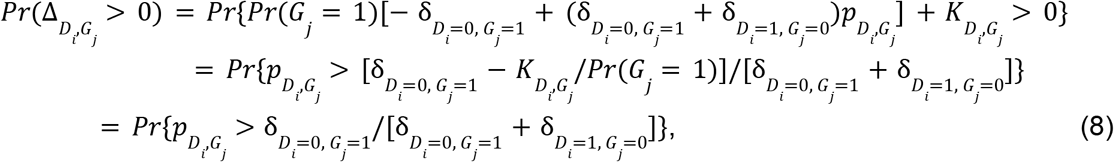

where the last line holds if 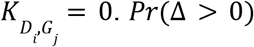 does not generally have a closed form, but can be calculated empirically from the sampling distributions of the 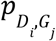 s. One can also derive a lower bound on the estimated 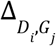 s that accounts for uncertainty by plugging in the fifth percentiles of the 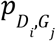 s in the uncertainty distributions in place of *Pr*(*D*_*i*_ = 1 | *G*_*j*_ = 1) in Equation (6). This fifth percentile represents a “near-worst case scenario” for the net utility in which the true disease penetrance is at the low end of its credible range.

## Data Availability

Data sharing is not applicable to this article as no new data were created or analyzed in this study.

## Code Availability

Our net utility calculations are available in an R Shiny app, freely accessible at https://janewliang.shinyapps.io/agg_utility. Code is available at https://github.com/janewliang/agg_utility.

## Acknowledgements

J.W.L. was supported by the National Cancer Institute at the National Institutes of Health (5T32CA009337). K.D.C. was supported by the National Human Genome Research Institute (K01HG009173).

## Author Contributions

J.W.L (conceptualization, formal analysis, methodology, software, visualization, writing-original draft, writing-review & editing). K.D.C. (conceptualization, writing-review & editing). R.C.G. (conceptualization, writing-review & editing). P.K (conceptualization, methodology, writing-review & editing).

