## Supplementary Material for "A framework evaluating the utility of multi-gene, multi-disease population-based panel testing that accounts for uncertainty in penetrance estimates"

**Supplemental Material**

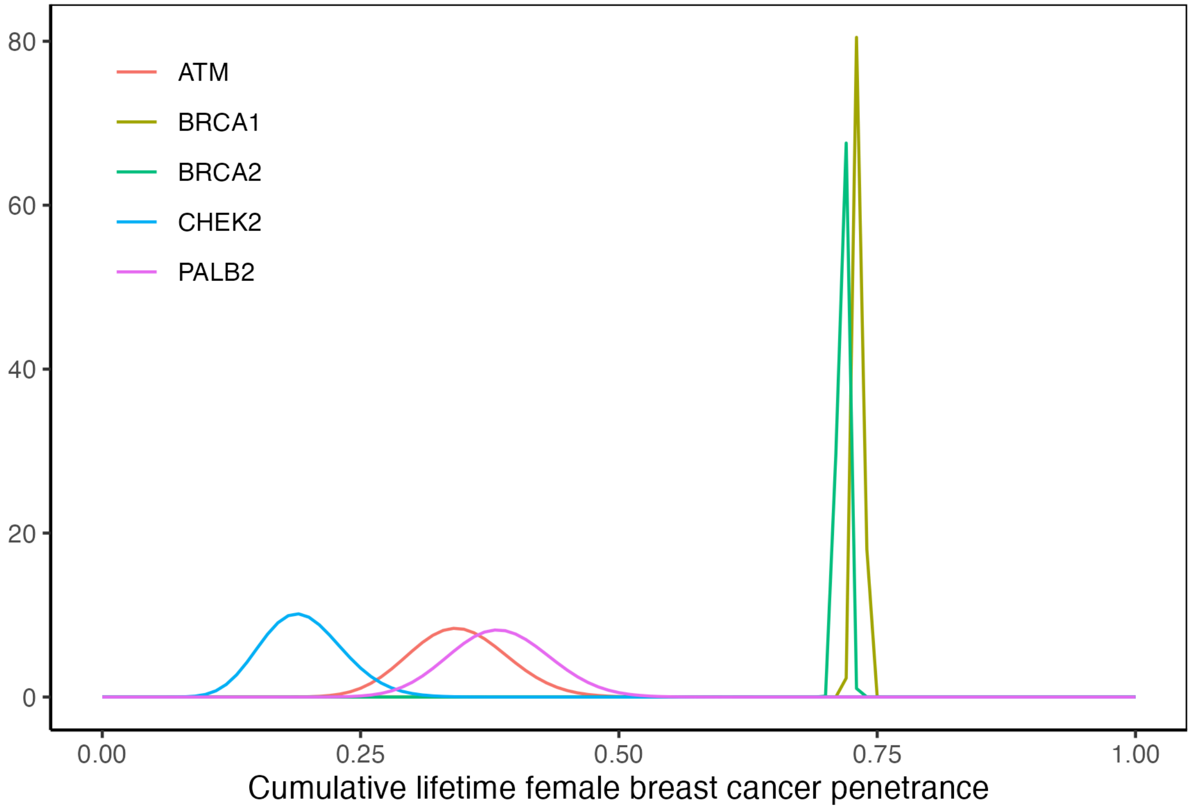


**Supplementary Figure 1**: Uncertainty distributions of the lifetime female breast cancer penetrance estimates for *ATM*, *BRCA1*, *BRCA2*, *CHEK2*, and *PALB2* carriers taken from a literature review (see text). For *BRCA1* and *BRCA2*, we used a precision of 10,000 to specify the uncertainty distribution, reflecting higher confidence about these estimates. For *ATM*, *CHEK2*, and *PALB2*, we used a precision of 100, because we have less certainty about these penetrances.

| | **0.025** | **0.05** | **0.10** | **0.50** | **0.90** | **0.95** | **0.975** |
|---:|---:|---:|---:|---:|---:|---:|---:|
| ***ATM*** | 0.256 | 0.269 | 0.285 | 0.344 | 0.407 | 0.425 | 0.441 |
| ***BRCA1*** | 0.723 | 0.725 | 0.726 | 0.732 | 0.738 | 0.739 | 0.741 |
| ***BRCA2*** | 0.708 | 0.709 | 0.711 | 0.717 | 0.722 | 0.724 | 0.725 |
| ***CHEK2*** | 0.124 | 0.133 | 0.145 | 0.193 | 0.246 | 0.263 | 0.277 |
| ***PALB2*** | 0.292 | 0.306 | 0.323 | 0.384 | 0.448 | 0.466 | 0.482 |

**Supplementary Table 1**: 2.5%, 5%, 10%, 50%, 90%, 95%, and 97.5% quantiles for the uncertainty distributions of the female breast cancer penetrances.

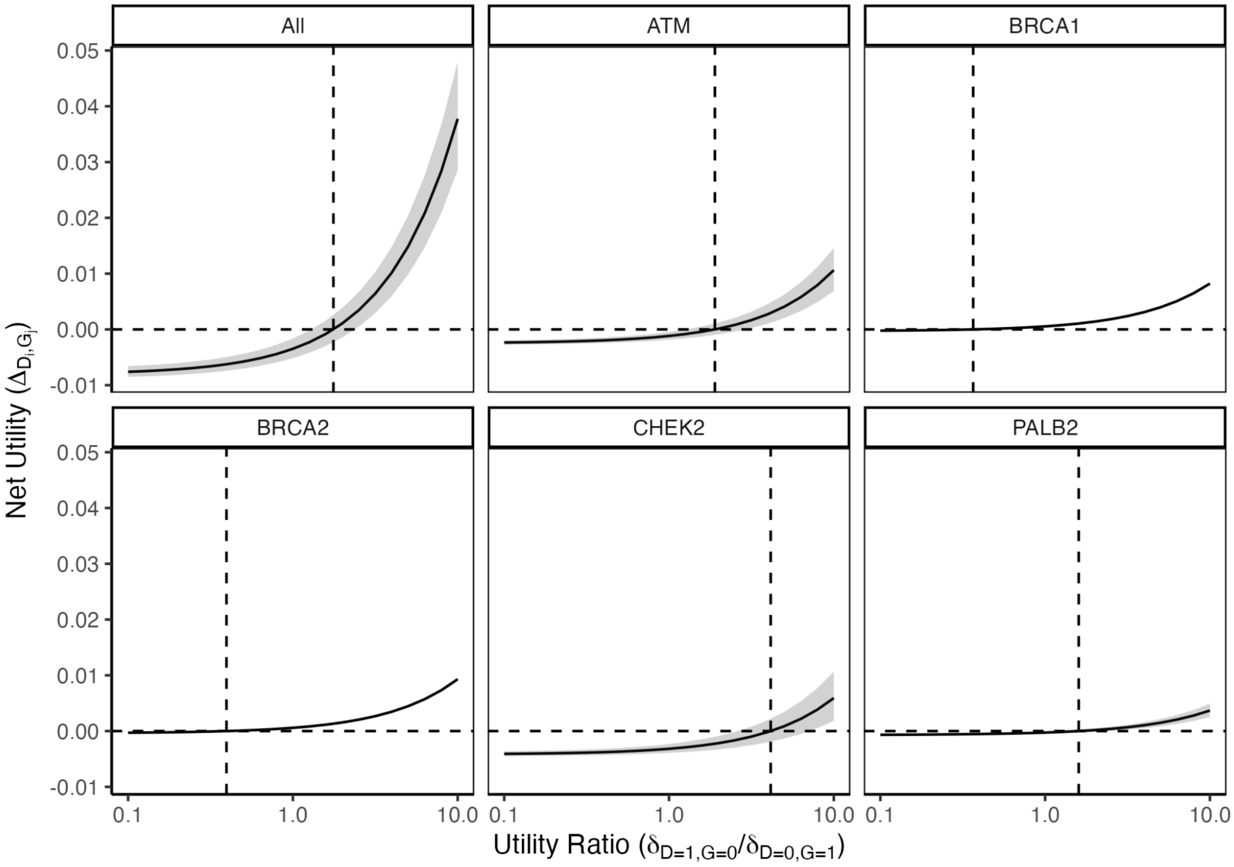


**Supplementary Figure 2**: Net utilities from the female breast cancer application plotted against the $\log_{10}$-scale ratio $\delta_{D=1,\,G=0}/\delta_{D=0,\,G=1}$ for each of the individual genes, as well as the aggregate $\Delta$ for all five genes. $\delta_{D=1,\,G=0}$ is allowed to vary from 0.1 to 10 in increments of $\log_{10}(0.1)$, while fixing $K = 0$ and $\delta_{D=0,\,G=1} = 1$. 95% credible intervals are shaded in gray and represent uncertainty contributed by the penetrance estimates taken from a literature review (see text). Dashed reference lines are drawn to indicate the value of the ratio at which the net utility changes from negative to positive.

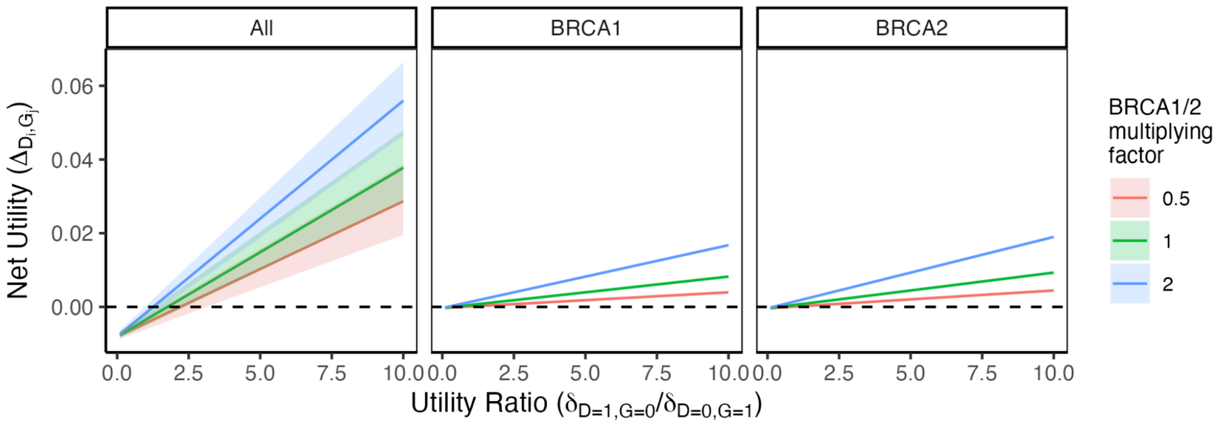


**Supplementary Figure 3:** Net utilities from the female breast cancer application plotted against the ratio $\delta_{D=1,\,G=0}/\delta_{D=0,\,G=1}$ for BRCA1, BRCA2, and the aggregate Δ for all five genes. 95% credible intervals are shaded. Dashed reference lines are drawn where the net utility is 0. We keep the inputs for the ATM, CHEK2, and PALB2 net utilities the same as in Figure 1 and Supplementary Figure 2, but multiply the utility ratios for BRCA1 and BRCA2 $\delta_{D_i=1,\,G_j=0}/\delta_{D_i=0,\,G_j=1} = \delta_{D=1,\,G=0}/\delta_{D=0,\,G=1}$ by factors of 0.5, 1, or 2. Because $\delta_{D_i=0,\,G_j=1}$ is normalized to 1, this is equivalent to considering a range of scenarios where $\delta_{D_i=1,\,G_j=0}$ for BRCA1 and BRCA2 is half, the same as, or twice that of the other 3 genes.

When the multiplying factor is 1, this reduces to the original analysis that assumes the same disutilities across all genes. When the factor is 0.5, $\delta_{D_i=1,\,G_j=0}/\delta_{D_i=0,\,G_j=1} = 0.5 * \delta_{D=1,\,G=0}/\delta_{D=0,\,G=1}$ for BRCA1 and BRCA2, thereby halving the penalty for G-D+ relative to G+D- (or doubling the penalty for G+D- relative to G-D+). The utility threshold at which the net utility is 0 is visibly higher for BRCA1, BRCA2, and the aggregate utility, indicating that we have restricted the spaces in which the BRCA1 and BRCA2 net utilities are positive, such that it would be beneficial to include the gene. These changes to the individual BRCA1 and BRCA2 net utility calculations then impact the aggregate utility. The interpretation when the factor is 2 and the utility threshold is visibly lower follows analogously.

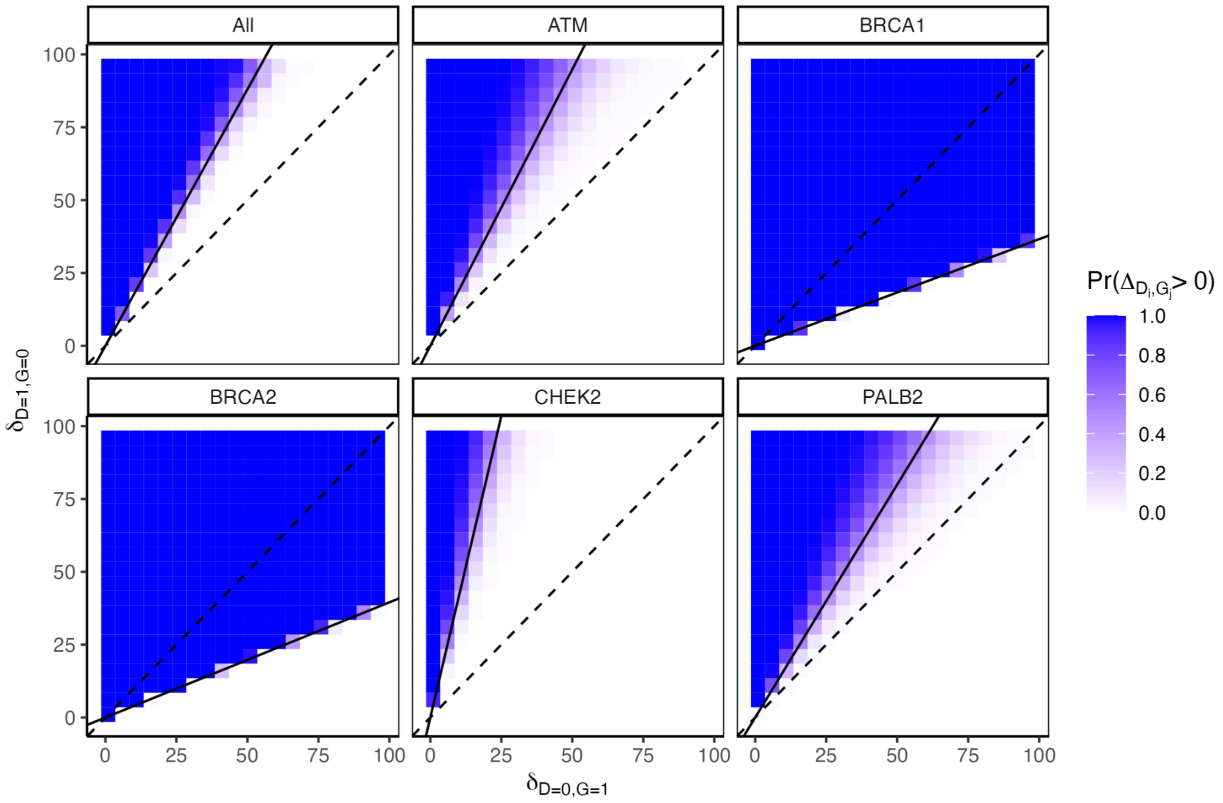


**Supplementary Figure 4**: Heatmaps of the probability of a positive net utility for the five individual female breast cancer tests and in aggregate ("All") while holding $K = 0$ and varying $\delta_{D=0,\, G=1}$ and $\delta_{D=1,\, G=0}$ from 1 to 100 in increments of 5. Probabilities near 1 are shaded blue and probabilities near 0 are shaded white. Utility thresholds based on the penetrance estimates are drawn as solid black lines. The dashed black lines have intercept 0 and slope 1, and correspond to cases where the individual testing positive for the gene but not developing the disease and vice versa have equal disutilities.

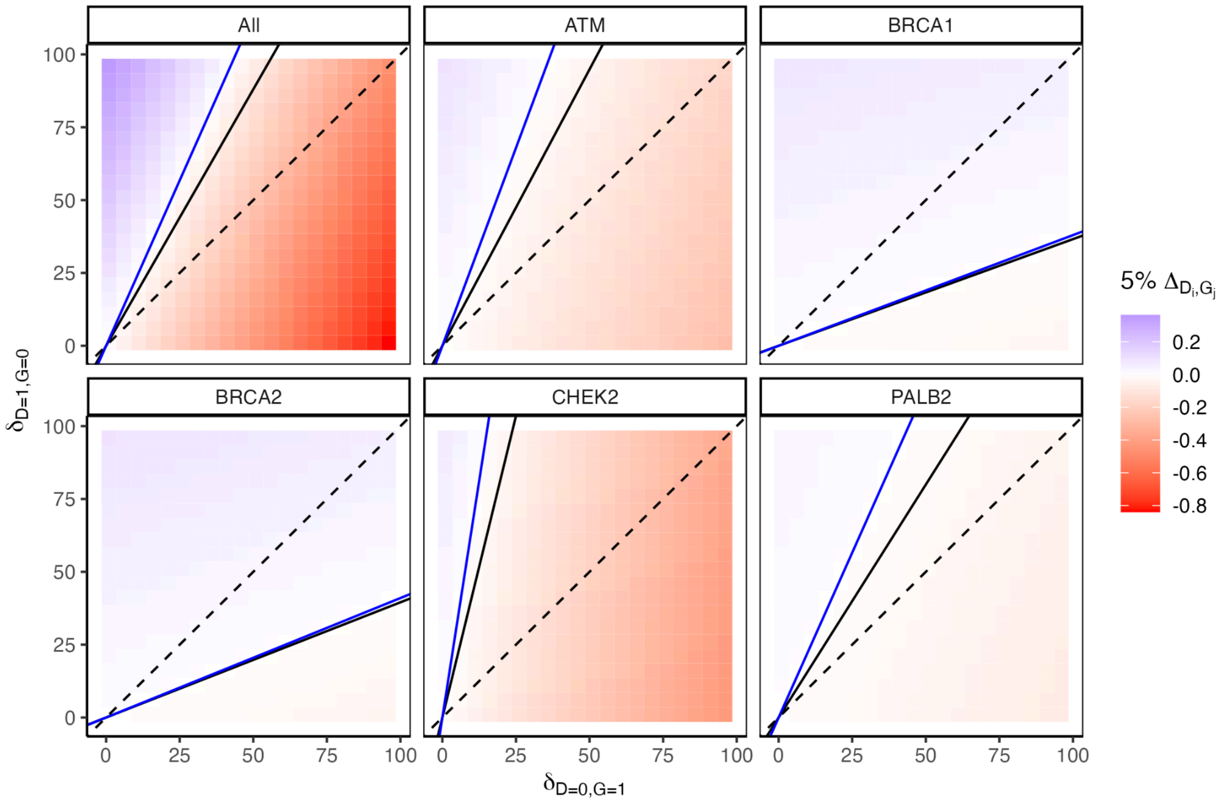


**Supplementary Figure 5**: Heatmaps of the five individual female breast cancer net utilities and the aggregate utility ("All"), based on the fifth percentiles of the penetrances' uncertainty distributions, while holding $K = 0$ and varying $\delta_{D=0,\,G=1}$ and $\delta_{D=1,\,G=0}$ from 1 to 100 in increments of 5. Positive fifth percentile net utilities are shaded blue and negative fifth percentile net utilities are shaded red. Utility thresholds based on the penetrance estimates are drawn as solid black lines. The dashed black lines have intercept 0 and slope 1, and correspond to utilities where the individual testing positive for the gene but not developing the disease and vice versa have equal disutilities. The solid blue lines indicate the utility thresholds based on the fifth percentiles of the penetrances' uncertainty distributions.

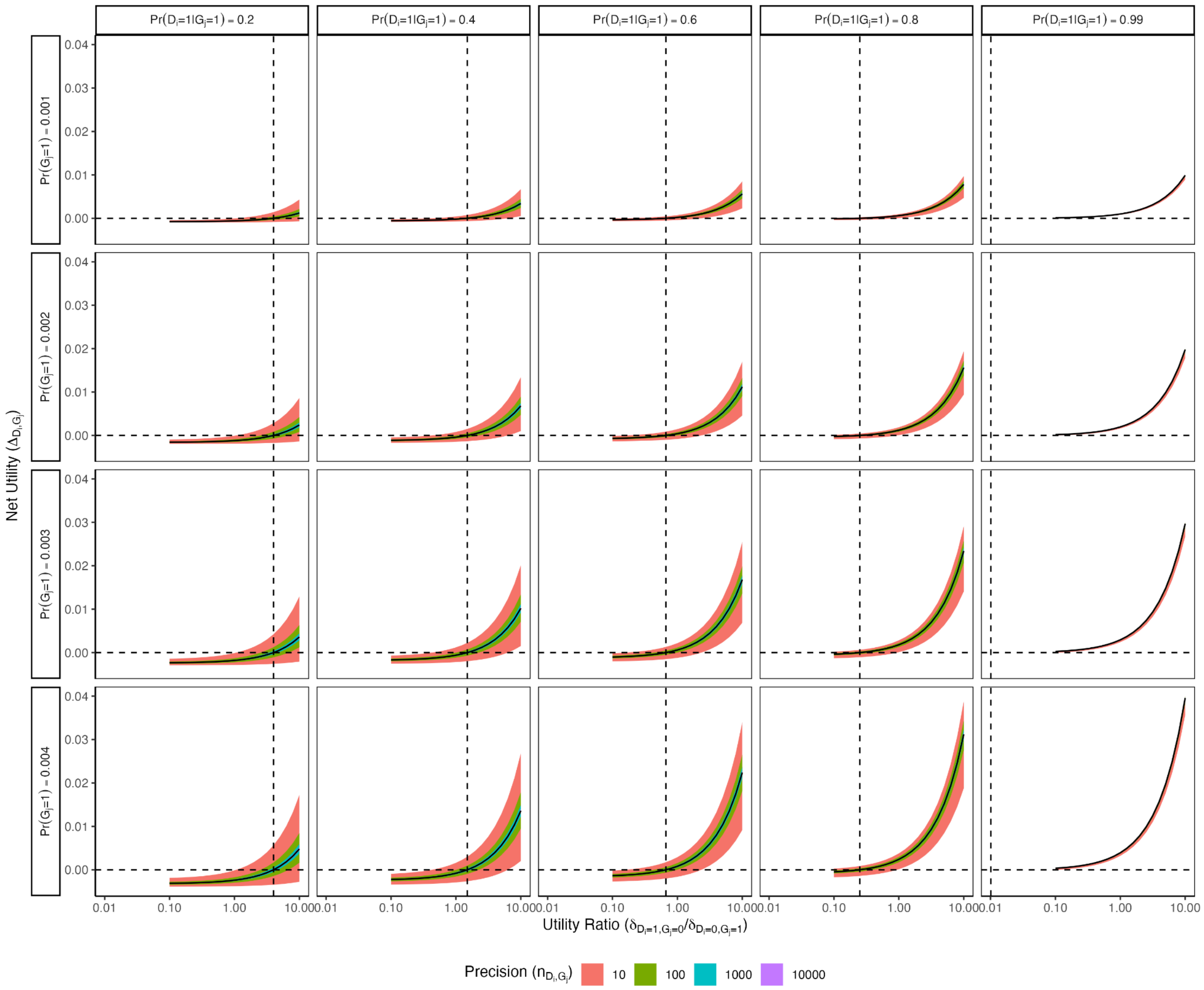


**Supplementary Figure 6**: Net utilities for a single gene and disease plotted against the $\log_{10}$ -scale utility ratio ($\delta_{D_i=1, G_j=0}/\delta_{D_i=0, G_j=1}$), while varying the G-D+ disutility, disease penetrance, carrier prevalence, and precision parameter. Test and G+D- disutilities are fixed at 0 and 1, respectively. The colored shading represents 95% credible intervals for different values of the precision. Dashed reference lines are drawn to indicate the utility threshold in each scenario.
